# Tissue embedding in a silica hydrogel for functional investigations

**DOI:** 10.1101/357962

**Authors:** Fabio Miazzi, Sabine Kaltofen, Jan E. Bello, Bill S. Hansson, Dieter Wicher

## Abstract

Embedding procedures can be challenging for delicate specimens. We present a method based on a sodium metasilicate (waterglass) gel to embed tissue samples for acute physiological studies. We show that the application of such a colloidal gel has minimal effects on the properties of buffered solutions and cell activity, allowing functional investigations on sensitive cells such as ciliated insect olfactory neurons.

## Introduction

Embedding conditions influence the mechanical and functional properties of cells and the outcome of physiological investigations (Chen et al., 2012). Moreover, they can be a source of thermal and osmotic stress that affects the cell physiology by inducing gene expression and changing metabolic activity to counteract non-optimal environmental conditions (Burg et al., 2007; Finan and Farshid, 2010; Richter et al., 2010). In order to minimize the negative effects of embedding media on samples, gelation agents should not induce changes in the physicochemical properties of the buffer, including pH, temperature and osmolality. Besides the well-known low melting temperature agarose (Ahrens et al., 2013; Kaufmann et al., 2012) and other natural source polymers, including alginate gels (de Vos et al., 2014), silica-based gels have been successfully used to encapsulate a variety of biological samples, ranging from enzymes to vertebrate cells (Avnir et al., 2006; Carturan et al., 2004; Nassif et al., 2002).

In particular, sodium metasilicate (Na_2_SiO_3_)-based aqueous solutions constitute an interesting case as they polymerize through a pH-driven condensation reaction (Pierre and Rigacci, 2011)

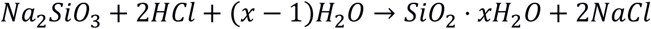

with salts and water as the only byproducts. Partial neutralization of basic Na_2_SiO_3_ solutions (pH ~12.5) to physiological (pH ~ 6.7-7.4) pH levels induce the gelation of colloidal silica particles through a chain of condensation reactions (Rao et al., 2011) (Fig. 1a). This means that such a process can be driven at room temperature and fixed pH levels in adequately buffered solutions. On this basis, we asked whether such a methodology could be used to embed tissue samples for acute physiological investigations via functional imaging techniques.

**Figure 1.**
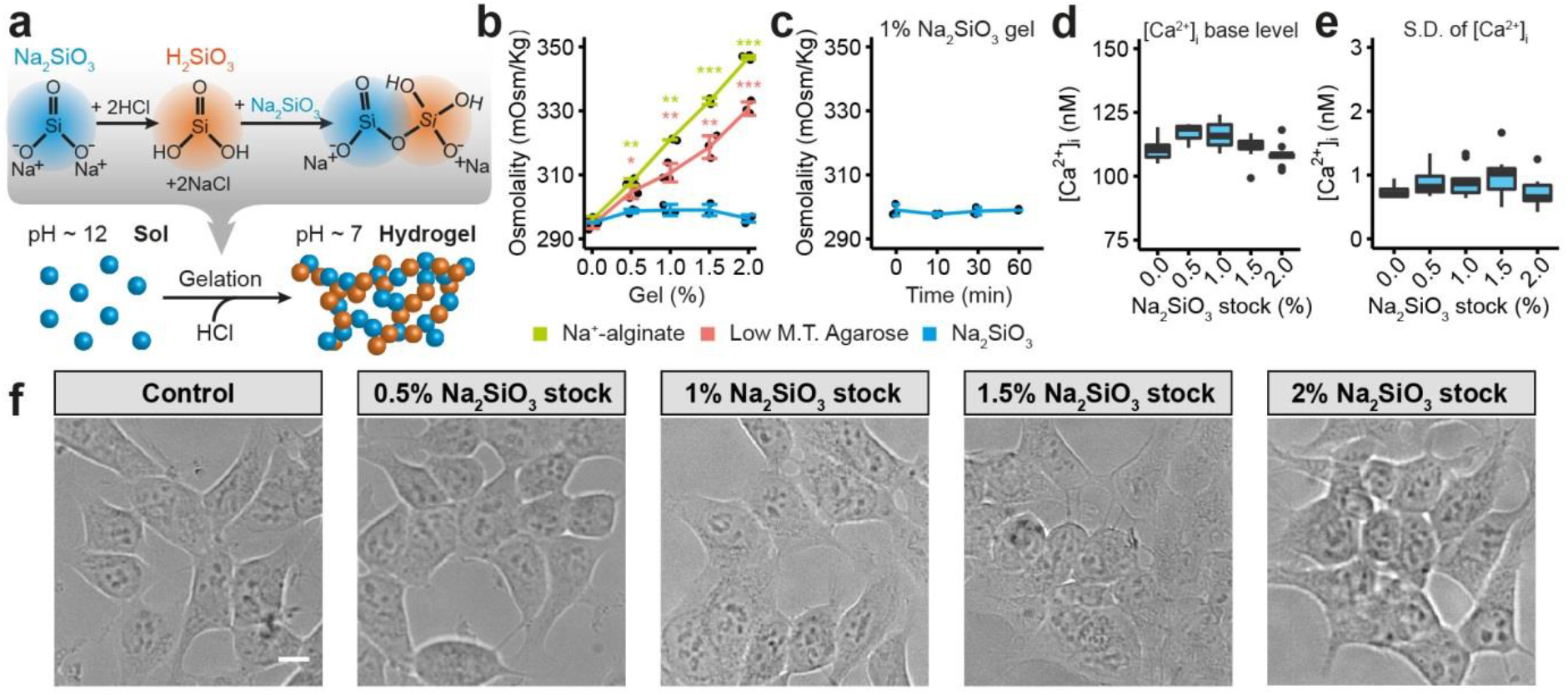
Sodium metasilicate (Na_2_SiO_3_) as an embedding agent for biological samples. (a) Schematic representation of the silicate gelation in pH buffered solutions. Partial neutralization of basic Na_2_SiO_3_ solutions with an acid produce silicic acid (H2SiO_3_) molecules that react with other Na_2_SiO_3_ or H_2_SiO_3_ molecules to form silicate colloidal particles. On a macroscopic scale condensation around these silicate particles lead to the formation of a silicate-based hydrogel (Rao et al., 2011). (b) Effect of common gelation agents and Na_2_SiO_3_ on the osmolality of a modified Hepes-HBSS buffered solution (see Methods section). Sodium alginate and low melting temperature (M. T.) agarose induce a concentration-dependent change in the medium osmolality that is significantly higher than Na_2_SiO_3_ colloidal particles. Na^2+^-alginate vs Na_2_SiO_3_: 0%: p = 0.4508; 0.5%, p = 0.002686; 1%, p = 0.00412; 1.5%, p = 0.00010528; 2%, p = 1.7388e-05. Agarose vs. Na_2_SiO_3_: 0%, p = 0.4508; 0.5%, p = 0.030430; 1%, p = 0.00718; 1.5%, p = 0.00355300; 2%, p = 1.0570e-04. Graph represents mean ± S.D., n = 3 for each concentration, two-tailed Welch’s t-tests with Holm’s multiple correction test. (c) Na_2_SiO_3_ polymerization does not affect the Hepes-HBSS buffer osmolality. Buffer osmolality was tested immediately or 10, 30 and 60 minutes after mixing the Na_2_SiO_3_, diluted HCl and Hepes-HBSS solutions (see Methods section). 0 vs 10 min: p = 0.9399, 0 vs. 30 min: p = 1, 0 vs. 60 min: p = 1. Graphs represents mean ± S.D., n = 3 for each concentration, two-tailed Welch’s t-tests with Holm’s multiple correction test. (d-e) Box and whiskers plots representing (d) the basal [Ca^2+^]_i_ levels at t = 0 and (e) the standard deviation (SD) of [Ca^2+^]i fluctuations of HEK293 cells incubated in buffer alone (0%) or containing increasing concentrations of Na_2_SiO_3_. n = 8 for 0% and 1.5% Na_2_SiO_3_ stock; n = 7 for 0.5% Na_2_SiO_3_ stock; n = 9 for 2% Na_2_SiO_3_ stock; n = 10 for 1% Na_2_SiO_3_ stock. (d) 0% vs 0.5%: p = 0.05596, 0% vs. 1%: p = 0.07995, 0% vs. 1.5%: p = 0.65640, 0% vs. 2%: p =0.65640. (e) 0% vs 0.5%: p = 0.21633, 0% vs. 1%: p = 0.34560, 0% vs. 1.5%: p = 0.19952, 0% vs. 2%: p = 0.48070. Two-tailed Wilkoxon-Mann-Whitney tests with Holm’s multiple test correction (see Methods section and Supplementary Fig. 2). Test statistic values, confidence intervals, degrees of freedom and uncorrected p values for panels (b-e) are reported in Supplementary Tables 1-3. * p < 0.05, ** p < 0.01, *** p < 0.001. (f) Morphology of HEK293 cells after incubation for one hour in presence of buffer alone (Control) or with increasing concentrations of sodium metasilicate (Na_2_SiO_3_). Cells tend to aggregate in clusters after incubation in presence of high Na_2_SiO_3_ concentrations (>1.5% Na_2_SiO_3_ stock, see online methods). Scale bar = 10 µm.

## Results and Discussion

We started by assessing the effects of the gelation process on the properties of pH-buffered solutions. We found that Na_2_SiO_3_ had significantly smaller effects on the buffer osmolality than other embedding media as agarose and sodium alginate (Fig. 1b) and this parameter was not affected by the silicate polymerization process (Fig. 1c, Supplementary Fig. 1), making sodium silicate hydrogels suitable for functional studies. We then explored the effects of Na_2_SiO_3_ on the dynamics of the morphology and the free intracellular calcium concentration ([Ca^2+^]_i_) of adherent HEK293 cells (Fig. 1d-f). Calcium imaging using the dye Fura2-AM demonstrated that Na_2_SiO_3_ did not affect the cell basal [Ca^2+^]_i_ (Fig. 1d) and did not induce variations of the [Ca^2+^]_i_ over time (Fig. 1e). Moreover, we observed changes in the cell morphology due to Na_2_SiO_3_ only at high concentrations (> 1.5% of a ≥ 27% SiO_2_ basis stock solution, see methods section), where cells tended to aggregate in clusters (Fig 1f). We can conclude that Na_2_SiO_3_ hydrogels are compatible with the embedding of living cells for functional studies.

We then explored the possibility of using the sodium metasilicate hydrogel system to embed biological samples employing the vinegar fly *Drosophila melanogaster* antennal tissue as a proof of concept. Insect antennae house olfactory sensory neurons (OSNs), expressing specialized olfactory receptors including odorant receptors (ORs) (Joseph and Carlson, 2015), which can be highly sensitive to air-borne compounds (Mansourian and Stensmyr, 2015). The OR-expressing olfactory cilia are encased in chitinous sensilla and isolated from the antennal endolymph by means of cell junctions between the OSNs and the surrounding supporting cells (Keil, 1999; Shanbhag et al., 2000), which makes it a very challenging task to access the OSNs and particularly the olfactory cilia. Moreover, in contrast to other species (Goetze et al., 2002; Stengl, 1994), as yet *Drosophila* OSNs have not been successfully cultured *in vitro*. In order to overcome such obstacles we tested whether Na_2_SiO_3_ can be used to embed dissociated antennal tissue samples for acute functional imaging investigations. By depositing the biological specimen on a glass coverslip previously cleaned to expose its hydrophilic surface, the gelation process could be used to anchor the sample to the glass support (Fig. 2a).

**Figure 2.**
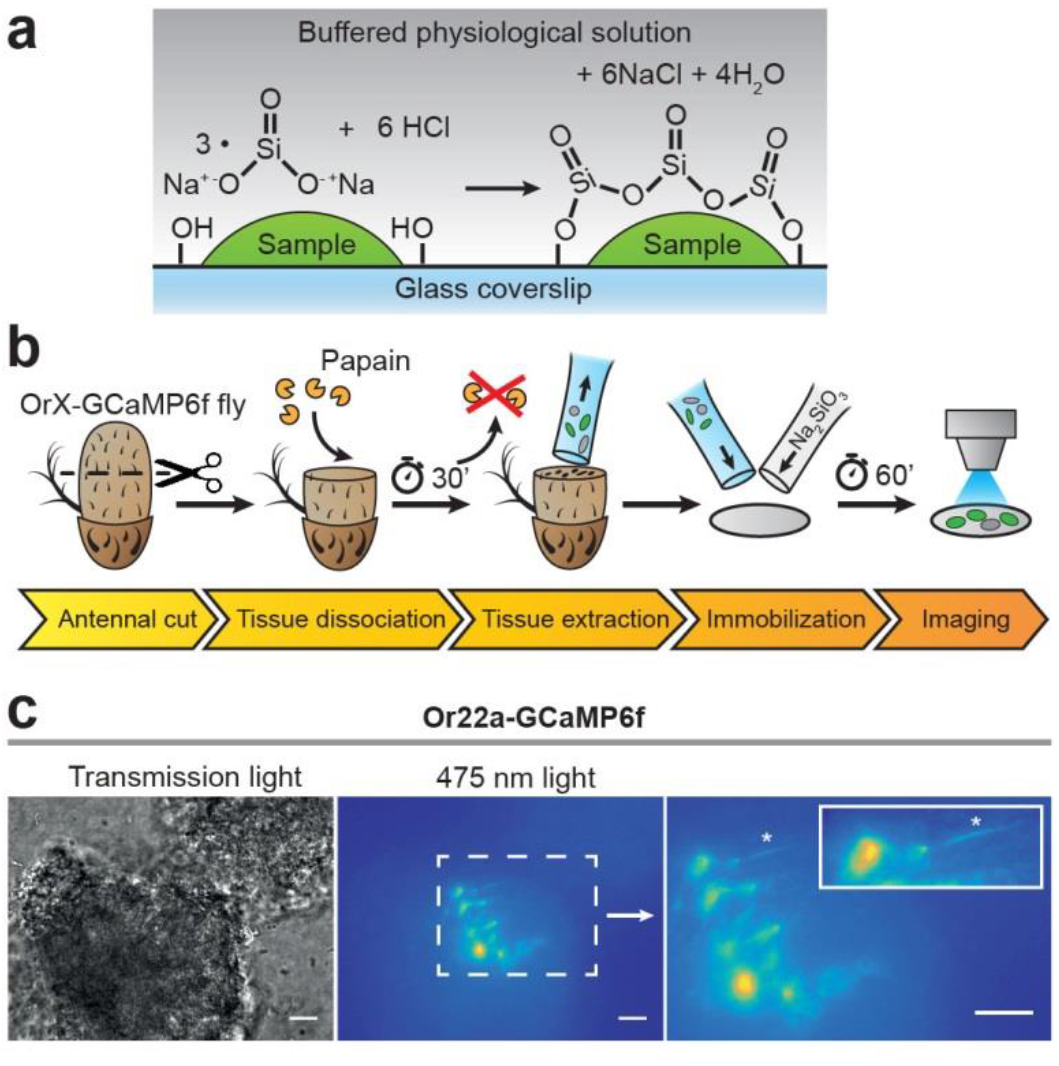
Embedding of dissociated *D. melanogaster* antennal tissue in a sodium metasilicate gel. (a) Scheme of sample embedding using Na_2_SiO_3_. When the sample is deposited on the hydrophilic glass surface, colloidal silica particles can covalently bind to it through a condensation reaction (top). In this way Na2SO3 can effectively encase the sample and avoid movement artifacts. (b) Schematic procedure for dissociation and embedding of vinegar fly antennal tissue (see Methods section). After fixing the antenna with a silicon-based curing medium, the funiculus was cut and incubated with a papain solution for 30 min. Then, the dissociated tissue was extracted with a silanized glass capillary and mixed with a modified *Drosophila* Schneider’s medium containing 0.972% of a Na_2_SiO_3_ stock solution (≥ 27% SiO_2_ basis). Ca2+ imaging was performed after a 60 min incubation time, to allow the Na_2_SiO_3_ gelation. (c) Example of dissociated and fixed antennal tissue under transmission (left) and 475 nm (center) light. The embedded tissue preserved the morphology of OSNs as they retained portions of their olfactory cilia (asterisk, right panel). Scale bar = 10 µm.

In order to assess the viability of the fly antennal olfactory sensory neurons (OSNs) and the extent of tissue damage following a transversal cut of the funiculus, we first set up an *in vivo* preparation to image OSNs after stimulations with airborne odors (Supplementary Fig. 3a-e). As a transversal cut of the funiculus did not induce *per se* a disruption of the antennal tissue sufficient to impair the ability of olfactory neurons to respond to odors and the vinegar fly OSNs retained their activity (Supplementary Fig. 3f-i, Supplementary Table 4), we developed a protocol to mildly digest and embed the fly antennal tissue. After funiculi from excised antennae were transversally cut, the antennal tissue was partially digested with papain – allowing the extraction of the digested content – and embedded on a clean glass coverslip using a modified Schneider’s medium containing 0.972% Na_2_SiO_3_ stock solution (Fig. 2b). This concentration of Na_2_SiO_3_ allowed us to reliably embed the tissue samples while having minimal effects on the cell physiology (Fig. 1). Following this procedure we obtained OSNs that retained their morphology, including the ciliated outer dendritic segment (Fig. 2c). We then asked whether the embedded neurons also retained their functional properties. We performed Ca^2+^ imaging from OSNs expressing the calcium indicator GCaMP6f (Chen et al., 2013) under the odorant receptor Or22a or the co-receptor Orco promoters and stimulated OSNs expressing Or22a or Orco with their respective agonists, ethyl hexanoate (Münch and Galizia, 2016) (Fig. 3a, Supplementary Video 1) and the synthetic compound VUAA1 (Jones et al., 2011) (Fig. 3b). After extraction of regions of interest (see Methods section), we calculated the changes in GCaMP6f fluorescence intensity with respect to the base level expressed in percent (% ΔF/F_0_). In both cases, OR agonists induced calcium responses in a concentration-dependent manner (Fig. 3c, d).

**Figure 3.**
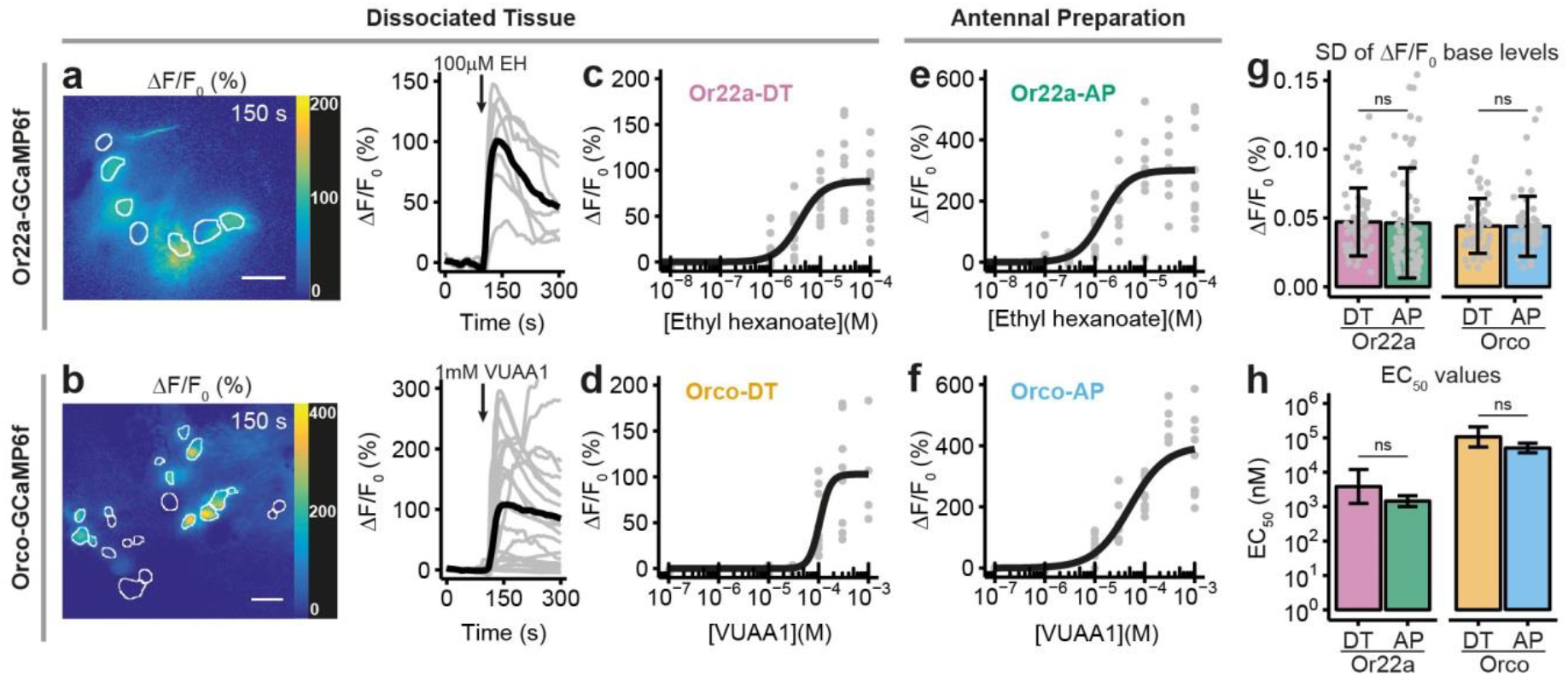
Comparison between Ca^2+^imaging from OSNs in sodium metasilicate-embedded and undissociated antennal tissue. (a-b) ΔF/F_0_ (%) from antennal dissociated tissue with (a) Or22a neurons (as in Figure 2c) and (b) Orco-expressing OSNs following the application of 100 μM ethyl hexanoate (EH) and VUAA1 respectively. White boundaries indicate regions of interest (ROIs) for quantitative analysis. Gray lines show the ΔF/F_0_ (%) of each ROI shown in the corresponding left panel; the black line represents the average taken for subsequent analysis (n = 1). Scale bar = 10 µm. (c-d) Concentration-response curves for Na_2_SiO_3_-embedded dissociated antennal tissue with Or22a (c, Or22a-DT) and Orco OSNs (d, Orco-DT) stimulated with ethyl hexanoate (8 ≤ n ≤ 10 for each concentration, 60 total data points) and VUAA1 (4 ≤ n ≤ 11 for each concentration, 49 total data points) respectively. (e-f) Concentration-response curves from Or22a (e) and Orco-expressing OSNs (f) from undissociated antennal tissue after stimulation with increasing concentrations of ethyl hexanoate (Or22a-AP) and VUAA1 (Orco-AP). 6 ≤ n ≤ 11 (e) and 5 ≤ n ≤ 11 (f) for each concentrati on, 71 and 60 total data points respectively. Parameters for curve fits are reported in the Supplementary Table 5. (g-h) Comparison between the ΔF/F_0_ standard deviation (SD) before stimulus application (g) and the half maximal effective concentrations (EC_50_) (h) of the OSNs from the undissociated antennal preparation (AP) and the dissociated tissue (DT) after stimulation of Or22a OSNs with ethyl hexanoate and Orco OSNs with VUAA1. (g) Or22a-AP: n = 71, Or22a-DN: n = 60, Or22a AP vs. DN: p = 0.9036. Orco-AP: n = 60, Orco-DN: n = 49, Orco AP vs. DN: p = 0.9816. Two-tailed Welch two sample t-tests (see the Methods section and Supplementary Fig. 6). Graphs represent mean ± S.D. Test statistic values, confidence intervals and degrees of freedom are reported in the Supplementary Table 6. (h) Or22a AP vs. DN: p = 0.1313. Orco AP vs. DN: p = 0.06676. Parameter comparison using the compParm function of the R (R Core Team, 2017) drc (Ritz et al., 2015) package. Statistic values are reported in the Supplementary Table 7. Graphs represent mean ± 95% C.I.

In order to determine if the OSN response profile was affected by the tissue dissociation process or Na_2_SiO_3_, we performed calcium imaging from excised antennae immediately after the funiculus cut (Fig. 3e-f and Supplementary Figure 4), and we compared the response profiles of Or22a and Orco-expressing OSNs from undigested and dissociated antennal tissue (Fig. 3g-h). Although the response profiles between the OSNs in undigested and dissociated tissues showed differences in the maximal intensity and time course of Ca^2+^ responses (Supplementary Fig. 5), we did not find significant differences in the fluctuation of the basal fluorescence levels (Fig. 3g) and in the EC_50_ of concentration-response curves for Or22a neurons stimulated with ethyl hexanoate and Orco-expressing neurons stimulated with VUAA1 (Fig. 3h). This suggests that the tissue dissociation and embedding procedures, at the Na_2_SiO_3_ concentration used, did not significantly affect the viability of OSNs as well as the diffusion of the OR agonists through the embedding medium.

In conclusion, we here demonstrate for the first time – to our knowledge – that sodium metasilicate hydrogels are excellent cell and tissue embedding agents for imaging experiments as they stabilize samples on uncoated glass surfaces, while retaining the function of neural cells without appreciable signs of cellular stress. The absence of toxic byproducts, the ability to form the gel at room temperature, together with the colloidal organization of silicate particles, which does not interfere with the osmolality of saline solutions during the gelation process, and its complete transparency make Na_2_SiO_3_ hydrogels an attractive choice, when embedding neural cell and tissue samples for physiological investigations.

## Materials and Methods

### Sodium metasilicate solution titration

A sodium metasilicate stock solution (≥ 27% SiO_2_ basis and ≥ 10% NaOH, Cat. Nr. 13729, Sigma-Aldrich, Steinheim, Germany) was first titrated in a 1:1000 dilution in double distilled water with 0.01 N HCl and pH changes were recoded with a pHmeter. The fit of the titration curve and the inflection point were calculated using CurTiPot (Gutz, I. G. R., CurTiPot – pH and Acid-Base Titration Curves: Analysis and Simulation software, version 4.2. http://www.iq.usp.br/gutz/Curtipot_.html). The stock sodium silicate solution had a [OH^−^] = 0.558 M.

### HEK293 cells culture and imaging

HEK293 cells (DSMZ Nr. ACC 305) were purchased from the Leibniz Institute DSMZ GmbH (Braunschweig, Germany) and grown in DMEM/F12 1:1 medium (Cat. Nr. 11320, Gibco, Life Technologies, Grand Island, NY, USA) supplied with 10% Fetal Bovine Serum at 37°C and 5% CO2. For experiments 80-90% confluent cells were dissociated by trypsinization and were subsequently cultured on poly-l-lysine (0.01%, Sigma-Aldrich, Steinheim, Germany) coated glass coverslips (12 mm diameter, Cat. Nr. P231.1, Carl Roth, Karlsruhe, Germany) at ~1 × 10^5^cells/well in a 24 wells plate with DMEM/F12 1:1 medium with 10% Fetal Bovine Serum at 37°C and 5% CO_2_. 24 hours post-plating, cells were incubated in Opti-MEM medium (Cat. Nr. 31985, Gibco), containing 5 µM Fura-2 acetoxymethyl ester (Molecular Probes, Invitrogen, Carlsbad, CA, USA) for 30 min at 37°C and 5% CO_2_. Then cells were washed twice in 26°C pre-warmed modified Hepes-HBSS (131.63 mM NaCl, 5.3 mM KCl, 0.5 mM MgCl_2_×6H_2_O, 1 mM EGTA, 0.2 mM CaCl_2_, 0.4 mM MgSO_4_, 0.44 mM KH_2_PO_4_, 4.17 mM NaHCO_3_, 0.34 mM Na_2_HPO_4_, 2 mM D-Glucose, 20 mM Hepes, pH = 7.4, osmolality = 296 mOsm/Kg) and 50 μl of modified Hepes-HBSS alone (Control) or with a sodium metasilicate (Na_2_SiO_3_) stock solution (≥ 27% SiO_2_ basis and ≥ 10% NaOH, Sigma-Aldrich) diluted 1:10 in double distilled water was applied on each coverslip (solutions: (i) 0.5% Na_2_SiO_3_ stock: 2.79 μl of HCl 0.05 M, 2.5 μl of Na_2_SiO_3_ diluted 1:10, 44.71 μl of modified Hepes-HBSS; (ii) 1% Na_2_SiO_3_ stock: 5.58 μl of HCl 0.05 M, 5 μl of Na_2_SiO_3_ diluted 1:10, 39.42 μl of modified Hepes-HBSS; (iii) 1.5% Na_2_SiO_3_ stock: 8.36 μl of HCl 0.05 M, 7.5 μl Na_2_SiO_3_ diluted 1:10, 34.14 μl of modified Hepes-HBSS; (iv) 2% Na_2_SiO_3_ stock: 11.15 μl of HCl 0.05 M, 10 μl Na_2_SiO_3_ diluted 1:10, 27.85 μl of modified Hepes-HBSS). Cells were incubated for one hour at 26°C in a humidified incubator to avoid desiccation. After incubation cells were kept throughout the experiment in standard extracellular solution (SES) containing 135 mM NaCl, 5 mM KCl, 1 mM CaCl_2_, 1 mM MgCl_2_×6H_2_O, 10 mM HEPES, 10 mM glucose (pH = 7.4; osmolality = 295 mOsmol/Kg). Excitation with 340 and 380 nm light was obtained using a monochromator (Polychrome V, TILL Photonics, Gräfelfing, Germany), coupled to an epifluorescence microscope (Axioskop FS, Zeiss, Jena, Germany) by means of a water immersion objective (LUMPFL40×W/IR/0.8; Olympus, Hamburg, Germany) and controlled by an imaging control unit (ICU, TILL Photonics). Emitted light was separated by a 400 nm dichroic mirror, filtered with a 420 nm long-pass filter and acquired by a cooled CCD camera (Sensicam, PCO Imaging, Kelheim, Germany) controlled by TILLVision 4.5 software (TILL Photonics). The exposure time was 150 ms per frame, with a temporal resolution of 0.2 Hz; in total each experiment lasted 50 s. The final image resolution was 640 × 480 pixels in a frame of 175 × 130 µm. Free intracellular Ca^2+^ concentration ([Ca^2+^]_i_) was calculated after background correction according to the equation:

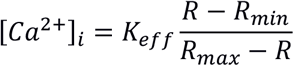

Where K_eff_ = 1.95 μM, R_min_ = 0.2 and R_max_ = 5.3. Regions of interest (ROIs) were selected using the built-in tools of TILL Vision and saved as comma-separated values (csv) files. Data analysis was performed using R (R Core Team, 2017) including add-on packages (Arnold, 2017; Auguie, 2017; Chang, 2014; Wickham, 2017). Non parametric statistics for HEK293 data analysis was used after evaluation of the pooled [Ca^2+^]_i_ distribution of all analyzed cells (ROIs). As the [Ca^2+^]_i_ distributions of the parameters analyzed in Figure 1b-c (Supplementary Fig. 2) show a long-tailed distribution, the median value for each independent replicate was used for subsequent non-parametric statistics (see the “Statistical methods” section below).

### Additional embedding media

Low melting temperature agarose (Cat. Nr. 6351, Roth) and sodium alginate (Cat. Nr. W201502, Sigma-Aldrich) in modified Hepes-HBSS at the desired concentration (% m/v) were incubated at 60 degrees until dissolved. Agarose solutions were cooled to 38 degrees before testing their osmolality, while sodium alginate solutions were cooled to room temperature and were tested without prior addition of calcium salts to induce their gelation. Osmolality of all solutions was tested with a Micro Osmometer 210 (Fiske, Norwood, MA, USA).

### Fly rearing and dissection procedure

*Drosophila melanogaster* with genotype *w; UAS-GCaMP6f; Orco-Gal4* and *w; UAS-GCaMP6f; Or22a-Gal4* were reared on conventional cornmeal agar medium under a 12h light: 12h dark cycle at 25°C. Orco-Gal4 and UAS-GCaMP6f were obtained from Bloomington Stock center (Nr. 42747 and Nr. 26818 respectively), while the Or22a-Gal4 line was kindly provided by Dr. Leslie Vosshall, Rockefeller University. Females between 4-8 days old were decapitated, the antennae were excised and fixed on a Sylgard-coated support (Sylgard 184, Dow Corning Corp. Midland, MI, USA) using a two-component silicon curing medium (Kwik-Sil, World Precision Instruments, Sarasota, FL, USA). The antennae were immersed in dissecting solution (Sicaeros et al., 2007) (Solution A: 137 mM NaCl, 5.4 mM KCl, 0.17 mM Na_2_HPO_4_, 0.22 mM KH2PO4. Solution B: 9.9 mM HEPES. For 500 ml of dissecting solution: 400 ml ultra-filtered water, 25 ml of Solution A, 14 ml of Solution B, 3.0 g (33.3 mM) D(+)-Glucose, 7.5g (43.8 mM) Sucrose. Brought to pH 6.7 with 1 N NaOH and to the final volume of 500 ml with ultra-filtered water) supplemented with 1 mM EDTA, 5 mM L-Cystein hydrochloride (Cat. Nr. C1276, Sigma-Aldrich) and, after equilibrating the pH to 6.7 with 1 N NaOH, 0.5 mg/ml Papain (Cat. Nr. 5125, Calbiochem, San Diego, CA, United States). Funiculi were cut between one third and one half of their length and incubated at 26°C for 30 minutes. After incubation, the dissecting solution was removed and the antennae were rinsed twice for 5 minutes at 26°C with Ca^2+^-free Ringer (130 mM NaCl, 4 mM MgCl_2_×6H_2_O, 36 mM sucrose, 5 mM HEPES, 5 mM KCl. Osmolality = 312 mOsm/Kg, pH = 6.7 adjusted with 1 N NaOH) supplemented with 1:75 cOmplete protease inhibitor cocktail (Cat. Nr. 04693116001, Roche, Basel, Switzerland) dissolved in a 100 mM PBS solution (50.93 mM Na_2_HPO_4_, 60.22 mM KH_2_PO_4_, 80.42 mM NaCl; pH = 6.7 adjusted with 1 N NaOH). A stock of Protease inhibitory solution was prepared by dissolving one tablet of cOmplete Protease Inhibitor Cocktail in 2 ml of 100 mM PBS. The solution was aliquoted and stored at −20°C for max 3 months. As an excessive concentration of cOmplete protease may cause cell permeabilization, the protease solution was added to the Ca^2+^-free Ringer in a 1:75 dilution, as suggested by the manufacturer.

### Glass coverslip preparation

Round glass coverslips (12 mm diameter, Cat. Nr. P231.1, Carl Roth, Karlsruhe, Germany) were cleaned by immersion in methanol (≥ 99.5%, Roth) and HCl (≥ 32%, Sigma-Aldrich) 1:1 for 30 minutes and after rinsing in double distilled water, by immersion in H_2_SO_4_ (95~97%, Sigma-Aldrich) for 30 minutes (Cras et al., 1999). Coverslips were then thoroughly rinsed and kept in methanol under N_2_; immediately before use, they were thoroughly washed in double distilled water and dried under N_2_.

### Glass capillaries and dissociated antennal tissue preparation

Borosilicate glass capillaries (0.86×150×80 mm, Cat. Nr. GB150-8P, Science Products, Hofheim, Germany) were pulled using a P-97 Micropipette puller (Sutter Instrument, Novato, CA, USA) and their tip was cut and fire polished in order to obtain holding micropipettes with an internal diameter ~0.4 mm. The capillary tip size was found to be critical in order to extract viable neurons. The inner diameter of the tip should be slightly larger than the width of the cut antenna when fixed in a vertical position. The capillaries were subsequently silanized by immersion for 10-15 seconds in 5% dimethildichlorosilan in toluene (Cat. Nr. 33065, Supelco, Sigma-Aldrich) and rinsed twice in toluene (≥99.5%, Sigma-Aldrich) and three times in methanol (≥ 99.5%, Sigma-Aldrich). The capillaries were then dried under N_2_ and heated at 200°C for two hours.

To embed the dissociated antennal tissue, a sodium metasilicate solution was prepared immediately before use by mixing 2.71 µl of HCl 0.05 M, 2.43 µl of sodium metasilicate solution (Cat. Nr. 13729, Sigma-Aldrich) diluted 1:10 in double distilled water and 13.86 µl of low Ca^2+^-Schneider’s medium (Cat. Nr. S9895, Sigma Aldrich, modified with 0.4 g EGTA (1 mM), 22.2 mg CaCl2 (free Ca^2+^ = 500 nM), 0.4 g NaHCO3, total volume: 1 liter, pH = 6.7 adjusted with 1 N NaOH; sterile filtered and kept at 4°C). After a 1 µl drop of Ca^2+^-free Ringer was deposited on a cleaned coverslip, the content of the treated antennae was gently sucked using a silanized capillary attached to a 2 µl micropipette. Usually ~0.5/1 µl of liquid was sucked together with each antenna. Within 10 minutes from mixing, the complete volume of the silicate gel solution was added to the coverslip and distributed uniformly. The total volume of liquid on a coverslip should be ~ 25 µl, for a final Na_2_SiO_3_ concentration equal to 0.972% of the Na_2_SiO_3_ (≥ 27% SiO_2_ basis) stock solution. Coverslips were incubated for one hour at 26°C in a high humidity environment, to avoid desiccation.

### Antennal preparation (without tissue dissociation)

For undissociated tissue antennal preparations, flies were decapitated, the antennae were excised and fixed on a Sylgard-coated support using the two-component silicon curing medium and immersed in *Drosophila* Ringer solution (130 mM NaCl, 5 mM KCl, 4 mM MgCl_2_×6H_2_O, 2 mM CaCl_2_, 36 mM sucrose, 5 mM Hepes, pH = 7.3, osmolality = 312 mOsm/Kg).

### Calcium imaging and data analysis from *D. melanogaster* olfactory neurons

Samples were immersed in *Drosophila* Ringer solution and imaged using the imaging setup described above with a LUMPFL 60x/0.90 water immersion objective (Olympus, Hamburg, Germany). Emitted light was separated by a 490 nm dichroic mirror and filtered with a 515 nm long-pass filter. GCaMP6f was excited with a 475 nm light for 50 ms per frame and a temporal resolution of 0.2 Hz. Stimuli consisted of 100 μl of ethyl hexanoate (99%, Sigma-Aldrich) or VUAA1 (N-(4-ethylphenyl)-2-((4-ethyl-5-(3-pyridinyl)-4H-1,2,4-triazol-3-yl)thio) acetamide, synthesized by the working group “Mass Spectrometry/Proteomics” of the Max Planck Institute for Chemical Ecology, Jena) at the required concentration. Ethyl hexanoate and VUAA1 working solutions were freshly prepared from 100 mM (or 500 mM, for final concentrations > 100 µM) stocks in DMSO, kept at −20°C. DMSO at a 1:1000 dilution in *Drosophila* Ringer was used as Control. Imaging data were exported as uncompressed tiff files and analyzed using custom scripts in Fiji-ImageJ2 (Rueden et al., 2017; Schindelin et al., 2012) where the regions of interest were selected using a semi-automatic procedure and the ΔF/F_0_ values were calculated after background, flat-field and movement correction. Statistical analysis was performed in R (R Core Team, 2017) using custom scripts including add-on packages (Arnold, 2017, 2017; Auguie, 2017; Chang, 2014; Ritz et al., 2015; Wickham, 2016). Parametric statistics for data analysis of the standard deviation of base level ∆F/F_0_ values for GCaMP6f (Fig. 3g) was used after evaluation of the pooled ∆F/F_0_ values distribution of all analyzed cells (ROIs) (Supplementary Fig. 6 and see the “Statistical methods” section below).

### Statistical methods

The appropriate statistics for data analysis on Figure 1d-e and Figure 3g was evaluated after assessment of the data distributions. For HEK293 cells relative frequency distributions – including all regions of interest (ROIs) selected for each treatment – were evaluated for the basal intracellular free Ca^2+^concentration ([Ca^2+^]_i_) at time t = 0 s (Supplementary Fig. 2a) and the standard deviation of the [Ca^2+^]_i_ during the 50 s time series imaging experiments (Supplementary Fig. 2b). In both cases, multiple treatments showed a distribution with a terminal long tail. As a consequence, extreme values of few cells can skew the mean value of each independent sample. For this reason, non-parametric statistics was applied to the HEK293 dataset: the median value between all ROIs was evaluated for each independent sample and comparisons between groups were performed using two tailed Wilcoxon-Mann-Whitney tests with Holm’s multiple test correction.

The same methodology was applied for the GCaMP6f ΔF/F_0_ data. The standard deviation of ΔF/F_0_ values before stimulus application was calculated for each ROI and the relative frequency distribution of values for each treatment was assessed (Supplementary Fig. 6). In this case, none of the treatments showed a terminal long tail, meaning that the number of ROIs with skewed high values – and consequently with the chance to skew the mean – was negligible. As a consequence, the mean value between all ROIs was selected for each independent measure and the difference between groups was evaluated using two tailed Welch’s t-tests.

### Functional calcium imaging from an *in vivo* antennal preparation

Flies with genotype *w, UAS-GCaMP3.0; +; Or22a-Gal4* (UAS-GCamP3.0 parental line was obtained from Bloomington Stock Center – Nr. 32234 –, Or22a-Gal4 line was kindly provided by Dr. Leslie Vosshall, Rockefeller University) were anesthetized in ice for 30 minutes and then placed into a truncated 1 ml pipette tip, leaving the head out of the tip and fixed using odor-free glue. The truncated tip was fixed using modeling clay on a custom Plexiglas mounting block (Strutz et al., 2012). Next, an antenna was fixed in vertical position by inserting it in a slit on a custom holder made of aluminum foil and a small plastic ring glued on a glass coverslip (18×24 mm, #0 thickness, Menzel-Gläser, Braunschweig, Germany). The arista was then glued on the top of the glass coverslip with odor-free glue and the funiculus was manually cut at around half of its length with a scalpel blade #22 (Fine Science Tools, Heidelberg, Germany). Immediately after cutting the antenna, a small glass coverslip (15×15 mm, #00 thickness, Menzel-Gläser) moistened with a very thin layer of halocarbon oil 700 (Cat. Nr. H8898, Sigma-Aldrich) was laid on the open funiculus to seal it. Both coverslips were fixed to each other and to the Plexiglas holder with odor-free glue to prevent movement artifacts. Imaging was performed using a BX51W1 widefield fluorescence microscope (Olympus) equipped with a DCLP490 dichroic mirror and a 60x/0.90 water immersion LUMPFL objective (Olympus). The objective was immersed in a drop of distilled water put on top of the coverslip sealing the funiculus. GCaMP3.0 stimulation with a 475 nm light and an exposure time of 50 ms was performed using a monochromator (Polychrome V, TILL Photonics). Emitted light was filtered with a LP515 filter and acquired at a 4 Hz frequency using a cooled CCD camera (Sensicam, PCO Imaging) controlled by TILLVision 4.5 software (TILL Photonics). Odor stimuli were sampled from the headspace of a 50 ml volume glass bottle (Schott, Jena, Germany) containing 2 ml of ethyl hexanoate (99% purity, Cat. Nr. 148962, Sigma-Aldrich) diluted in mineral oil to a 10-2 dilution. A stimulus controller (Syntech, Kirchzarten, Germany) was used to deliver for 2 s the bottle headspace into a charcoal-filtered and humidified constant air flow (0.5 m/s) in a Teflon tube, which inlet was positioned 5-10 mm from the recorded antenna. The stimulus was delivered at 10 s from the start of the recording; the total duration of the experiment was 35s. The response magnitude was calculated for each frame as the average ΔF/F_0_ and expressed in percentage after background subtraction. Regions of interest (ROIs) were selected using the built-in tools of TILLVision 4.5 (Supplementary Fig. 3d-e) and F_0_ was estimated as the mean fluorescence level calculated for each selected ROI as the average intensity from 3 to 5.25 s of the recording. Data analysis was performed using Prism 4 software (GraphPad Software Inc., La Jolla, CA, USA).

## Data availability statement

All data analyzed during this study and the custom scripts used for data analysis are available from the corresponding author on reasonable request.

## Acknowledgements

The authors thank Silke Trautheim for help rearing the fly lines and Aleš Svatoš and Jerrit Weißflog for the synthesis of VUAA1. This project received funding from the International Max Planck Research School (FM), the European Union’s Horizon 2020 research and innovation program under the Grant Agreement No. 662629 (FM), the Alexander von Humboldt Foundation (JEB) and the Max Planck Society (DW, SK, BHS). Stocks obtained from the Bloomington Drosophila Stock Center (NIH P40OD018537) were used in this study.

## Author contributions

DW, FM and BSH formulated the ideas for the study, FM and DW designed the study, JEB contributed to study design, FM and SK performed the experiments, FM analyzed the data and wrote the manuscript. All authors contributed to the final version of the manuscript.

